# Comparing tonic and phasic calcium in the dendrites of vulnerable midbrain neurons

**DOI:** 10.1101/2023.08.28.555184

**Authors:** Rita Yu-Tzu Chen, Rebekah C. Evans

**Affiliations:** Department of Neuroscience, Georgetown University Medical Center, Washington DC

## Abstract

Several midbrain nuclei degenerate in Parkinson’s Disease (PD). Many of these nuclei share the common characteristics that are thought to contribute to their selective vulnerability, including pacemaking activity and high levels of calcium influx. In addition to the well-characterized dopaminergic neurons of the substantia nigra pars compacta (SNc), the cholinergic neurons of the pedunculopontine nucleus (PPN) also degenerate in PD. It is well established that the low-threshold L-type calcium current is a main contributor to tonic calcium in SNc dopaminergic neurons and is hypothesized to contribute to their selective vulnerability. However, it is not yet clear whether the vulnerable PPN cholinergic neurons share this property. Therefore, we used two-photon dendritic calcium imaging and whole-cell electrophysiology to evaluate the role of L-type calcium channels in the tonic and phasic activity of PPN neurons and the corresponding dendritic calcium signal and directly compare these characteristics to SNc neurons. We found that blocking L-type channels reduces tonic firing rate and dendritic calcium levels in SNc neurons. By contrast, the calcium load in PPN neurons during pacemaking did not depend on L-type channels. However, we find that blocking L-type channels reduces phasic calcium influx in PPN dendrites. Together, these findings show that L-type calcium channels play different roles in the activity of SNc and PPN neurons, and suggest that low-threshold L-type channels are not responsible for tonic calcium levels in PPN cholinergic neurons and are therefore not likely to be a source of selective vulnerability in these cells.

## Introduction

Parkinson’s disease (PD) is a disabling neurodegenerative disease characterized by motor deficits such as bradykinesia and rigidity, as well as cognitive decline (Poewe et al., 2017). The cardinal symptoms of PD have been attributed to loss of dopaminergic neurons in the substantia nigra pars compacta (SNc), but several other brainstem nuclei also degenerate (Braak et al., 2004; Giguère et al., 2018). Of particular interest, ∼30-60% of cholinergic pedunculopontine nucleus (PPN) neurons are lost in PD patients (Giguère et al., 2018; Jellinger, 1988; Rinne et al., 2008; Sébille et al., 2019). PPN cholinergic neurons directly innervate motor structures in the basal ganglia and lower brainstem (Mena-Segovia and Bolam, 2017), and their degeneration may contribute to gait and balance impairments (Chambers et al., 2021; Grabli et al., 2013; Karachi et al., 2010; Rinne et al., 2008). However, it is not known why PPN cholinergic neurons and SNc dopaminergic neurons selectively degenerate in PD.

The factors that make some neurons vulnerable to degeneration while others are resilient have been elusive. While there are multiple hypotheses for selective vulnerability (Giguère et al., 2018; Gonzalez-Rodriguez et al., 2020), one of particular therapeutic interest is that tonic pacemaking and the accompanying calcium (Ca^2+^) influx increases neural vulnerability. The pacemaking activity of SNc dopaminergic neurons is well characterized (Grace and Onn, 1989; Johnson et al., 1992). It is accompanied by large somatodendritic Ca^2+^ oscillations primarily mediated by low-threshold L-type Ca^2+^ channels (Chan et al., 2007; Guzman et al., 2009; Hage and Khaliq, 2015; Shin et al., 2022). Because this chronic Ca^2+^ load adds to metabolic cost (Surmeier et al., 2010) and contributes to mitochondrial oxidative stress (Dryanovski et al., 2013; Guzman et al., 2010), L-type Ca^2+^ channel blockers have been investigated as treatment to slow the progress of PD in animal models (Liss and Striessnig, 2019; Ortner, 2021) and in clinical trials (Parkinson Study Group STEADY-PD III Investigators, 2020; Surmeier et al., 2022).

PPN cholinergic neurons, like SNc dopaminergic neurons, spontaneously fire action potentials in the absence of excitatory synaptic input (Takakusaki and Kitai, 1997). However, it is not known whether this tonic pacemaking contributes to their vulnerability in PD or whether they share pacemaking mechanisms with the vulnerable SNc dopaminergic neurons. Previous studies indicate that PPN cholinergic neurons undergo Ca^2+^-dependent membrane potential oscillations (Hyde et al., 2013; Kezunovic et al., 2011; Takakusaki and Kitai, 1997), but dendritic Ca^2+^ activity during pacemaking has not yet been measured. To determine whether PPN cholinergic neurons share the same pacemaking and Ca^2+^ influx mechanisms as SNc dopaminergic neurons, we used whole-cell patch clamp with simultaneous two-photon Ca^2+^ imaging to measure dendritic Ca^2+^ during tonic and phasic action potential firing. We find that PPN cholinergic neurons exhibit pacemaking Ca^2+^ that is highly associated with action potential spiking, but unlike SNc neurons, this tonic Ca^2+^ is not mediated by L-type channels. However, we find that L-type channels contribute to phasic firing-induced Ca^2+^ entry in PPN neurons, suggesting that they express high-threshold L-type channels. Therefore, these findings reveal that L-type channels play different roles in the regulation of tonic and phasic Ca^2+^ dynamics in PPN and SNc neurons, and show that PPN cholinergic neurons do not rely on low-threshold L-type channels for spontaneous pacemaking or tonic Ca^2+^ levels.

## Methods

### Animals

All animal procedures were approved by the Georgetown University Medical Center Institutional Animal Care and Use Committee (IACUC). ChAT-Cre (strain #031661) and Ai9/tdTomato (strain #007909) mice on the C57BL/6J background were purchased from Jackson Laboratory and crossed to produce ChAT-Cre/TdTomato mice. WT C57BL/6J (strain #000664) mice were purchased from Jackson Laboratory and bred in the Georgetown University Department of Comparative Medicine animal facility. Mice were group-housed with same-sex littermates when possible and had *ad libitum* access to food and water in a 12-hr light/12-hr dark cycle room.

### Slice preparation

Horizontal brain slices (200 μm) were prepared from adult ChAT-Cre/tdTomato or WT mice of either sex (>2 months old, average age = 110 ± 5 days, range = 70-140 days). Mice were anesthetized with isoflurane and transcardially perfused with ice-cold slicing solution containing (in mM): 198 glycerol, 25 NaHCO_3_, 2.5 KCl, 1.2 NaH_2_PO_4_, 20 HEPES, 10 glucose, 10 MgCl_2_, 0.5 CaCl_2_ (bubbled with 95% O_2_/5% CO_2_, osmolarity = 310-320 mmol/kg). Brains were then quickly extracted and slices were prepared using Leica VT1200S vibratome in the same slicing solution. The slices were transferred to and incubated for 30 min in a heated (34 °C) modified ACSF containing (in mM): 92 NaCl, 30 NaHCO_3_, 2.5 KCl, 1.2 NaH_2_PO_4_, 20 HEPES, 35 glucose, 2 MgCl_2_, 2 CaCl_2_, 5 Na-ascorbate, 3 Na-pyruvate, 2 thiourea (bubbled with 95% O_2_/5% CO_2_, osmolarity = 300-310 mmol/kg). After incubation, slices were moved to room temperature for at least an additional 30 min before recording.

### Electrophysiology

Slice were transferred to a recording chamber with constant perfusion of warm (30-34 °C) ACSF containing (in mM): 125 NaCl, 25 NaHCO_3_, 3.5 KCl, 1.25 NaH_2_PO_4_, 10 glucose, 1 MgCl_2_, 2 CaCl_2_ (bubbled with 95% O_2_/5% CO_2_, osmolarity = 295-310 mmol/kg). The pedunculopontine nucleus (PPN) was identified by tdTomato fluorescence in the ChAT+ neurons and its relative location to the superior cerebellar peduncle and the laterodorsal tegmental nucleus. Similar numbers of ChAT+ cells from the pars dissipata of rostral PPN and the pars compacta of caudal PPN were sampled for each drug treatment. Dopaminergic neurons of the substantia nigra pars compacta (SNc) were identified by their large soma size and relative location to the medial terminal nucleus of the accessory optic tract, as well as electrophysiological properties including slow pacemaking (1-8 Hz) and prominent voltage sag in response to hyperpolarizing current injection.

Whole-cell current-clamp recordings were made with a MultiClamp 700B amplifier and digitized with Digidata 1550B (Molecular Devices). Patch pipettes of tip resistance 2-6 MΩ were pulled from filamented borosilicate glass and filled with the intracellular solution containing (in mM): 122 KMeSO_3_, 9 NaCl, 9 HEPES, 1.8 MgCl_2_, 14 phosphocreatine, 4 Mg-ATP, 0.3 Tris-GTP, 0.05 Alexa Fluor 594, 0.3 Fluo-5F (pH = 7.35 with KOH; osmolarity = 290-300 mmol/kg). Current-clamp recordings were bridge balanced and liquid junction potential (−8 mV) was not corrected. For analysis of spontaneous tonic firing properties, only cells that were actively pacemaking were included.

### Two-photon calcium imaging

The majority of PPN cholinergic neurons exhibited slow pacemaking once the intracellular solution dialyzed the cells. A minority of PPN neurons (15%, 4 out of 26 cells) remained hyperpolarized and quiescent. During Ca^2+^ imaging experiments, which was initiated ≥15 min after whole-cell break-in, these 4 quiescent cells were injected with a constant amount of depolarizing current to maintain stable pacemaking (ranging from +30 to +120 pA). Ca^2+^ imaging was performed using previously published procedures (Evans et al., 2017). Cells were imaged on a two-photon microscope (Bruker) with a Mai Tai ultrafast Ti:sapphire laser (Spectra-Physics) tuned to 810 nm, which activates both Alexa Fluor 594 and Fluo-5F (Sabatini et al., 2002) but not tdTomato (Drobizhev et al., 2011). Linescans (2 ms lineperiod, 12 μs dwell time, 2 s total each scan) of the somatodendritic regions were taken at 512×512 pixels resolution using a 40×/0.8 NA objective (Olympus). Fluorescence signals were split into red and green channels by a 575 nm dichroic long-pass mirror and passed through 607/45 nm and 525/70 nm filters before being detected by multi-alkali photomultiplier tubes (Bruker). To visualize cell morphology, Z-stacks (1 μm step, 2 or 4 μs dwell time) of each recorded cell were taken at 512×512 pixels resolution after Ca^2+^ imaging experiments. Ca^2+^ signals were quantified as the ratio of green to red fluorescence (G/R), normalized to the ratio of saturated green to red signals (Gs/R), which were measured daily by placing a pipette filled with intracellular solution plus saturating Ca^2+^ (2 mM) at the surface of the brain slice.

### Drugs

All salts were from Sigma-Aldrich. Alexa Fluor 594 (Invitrogen/Life Technologies), Fluo-5F (Invitrogen/Life Technologies), and tetrodotoxin-citrate (Hello Bio) were dissolved in deionized water as concentrated stocks. Nifedipine (Tocris) was dissolved in DMSO, stored frozen and protected from light, and thawed only once on the day of use. The final concentration of DMSO in the ACSF was 0.03% (v/v) for nifedipine treatment and 0.05% (v/v) for control. After baseline measurements were taken, TTX was perfused in bath for 5 min and nifedipine for 8 min before post-drug measurements.

### Data analysis

Cell morphology and Ca^2+^ imaging data were quantified using ImageJ to determine distances or fluorescence signal intensities. All numerical data including electrophysiological traces were analyzed and graphed in Igor Pro (WaveMetrics). The Mann-Whitney-Wilcoxon test was used to compare two unpaired samples, while the Wilcoxon signed-rank test was used to compare two paired samples. The Pearson correlation coefficient (r) was used to determine the significance of linear correlation data. All results in text are reported as mean ± SEM. Box plots show medians as the middle line, 25^th^ and 75^th^ percentiles as the bottom and top of the box, and 10^th^ and 90^th^ percentiles as the whiskers. Measurements from the same cell at before and after conditions are linked by a line between the markers in box plots and analyzed as paired data. In all figures, one asterisk (*) denotes a statistical significance level of P value <0.05, two asterisks (**) P value <0.01, three asterisks (***) P value <0.001, and four asterisks (****) P value <0.0001.

## Results

### SNc dopaminergic and PPN cholinergic neurons differ in tonic firing properties

The boundary of the PPN is best defined by its choline acetyltransferase (ChAT) positive cholinergic neurons (Rye et al., 1987). In this study, we identified PPN cholinergic neurons by red fluorescence in brain slices prepared from ChAT-tdTomato transgenic mice. Dopaminergic neurons of the SNc were identified in WT or ChAT-tdTomato mice by using anatomical location and electrophysiological characteristics. SNc dopaminergic neurons and PPN cholinergic neurons both exhibit slow, spontaneous pacemaking activity. Using two-photon laser scanning microscopy combined with whole-cell patch-clamp, we visualized the dendritic morphology and recorded action potentials from those two neuronal populations by filling the cells with the red fluorescent dye Alexa Fluor 594.

Morphologically, SNc dopaminergic and PPN cholinergic neurons differ in soma shape and the orientations of primary dendrites. The majority of SNc neurons recorded have large, spindle-shaped somas (75%, 6 out of 8 cells) and primary dendrites that extend parallel to the tapered ends of the soma (Fig. 1A). The average soma dimensions of SNc neurons estimated from the Z-stacks are 25.38 ± 1.73 μm in length and 13.89 ± 0.44 μm in width. In comparison, most PPN neurons (85%, 23 out of 27 cells) have large multipolar somas with 2-4 primary dendrites coming off in multiple directions (Fig. 1B), with a minority (15%, 4 out of 27 cells) having spindle-shaped somas. These cholinergic neurons have an average length of 26.96 ± 0.96 μm and width of 16.77 ± 0.39 μm. The membrane capacitance (Cm) of SNc and PPN neurons did not differ significantly (SNc Cm: 79.18 ± 5.19 pF, n = 7; PPN Cm: 81.23 ± 3.44 pF, n = 26; Mann-Whitney-Wilcoxon test, p = 0.8803; Fig. 1F). However, PPN neurons exhibited higher input resistance (Ri) than SNc neurons (PPN Ri: 369.7 ± 26.7 MΩ, n = 26; SNc Ri: 170.3 ± 23.0 MΩ, n = 7; Mann-Whitney-Wilcoxon test, p = 0.0004; Fig. 1G).

**Figure 1.**
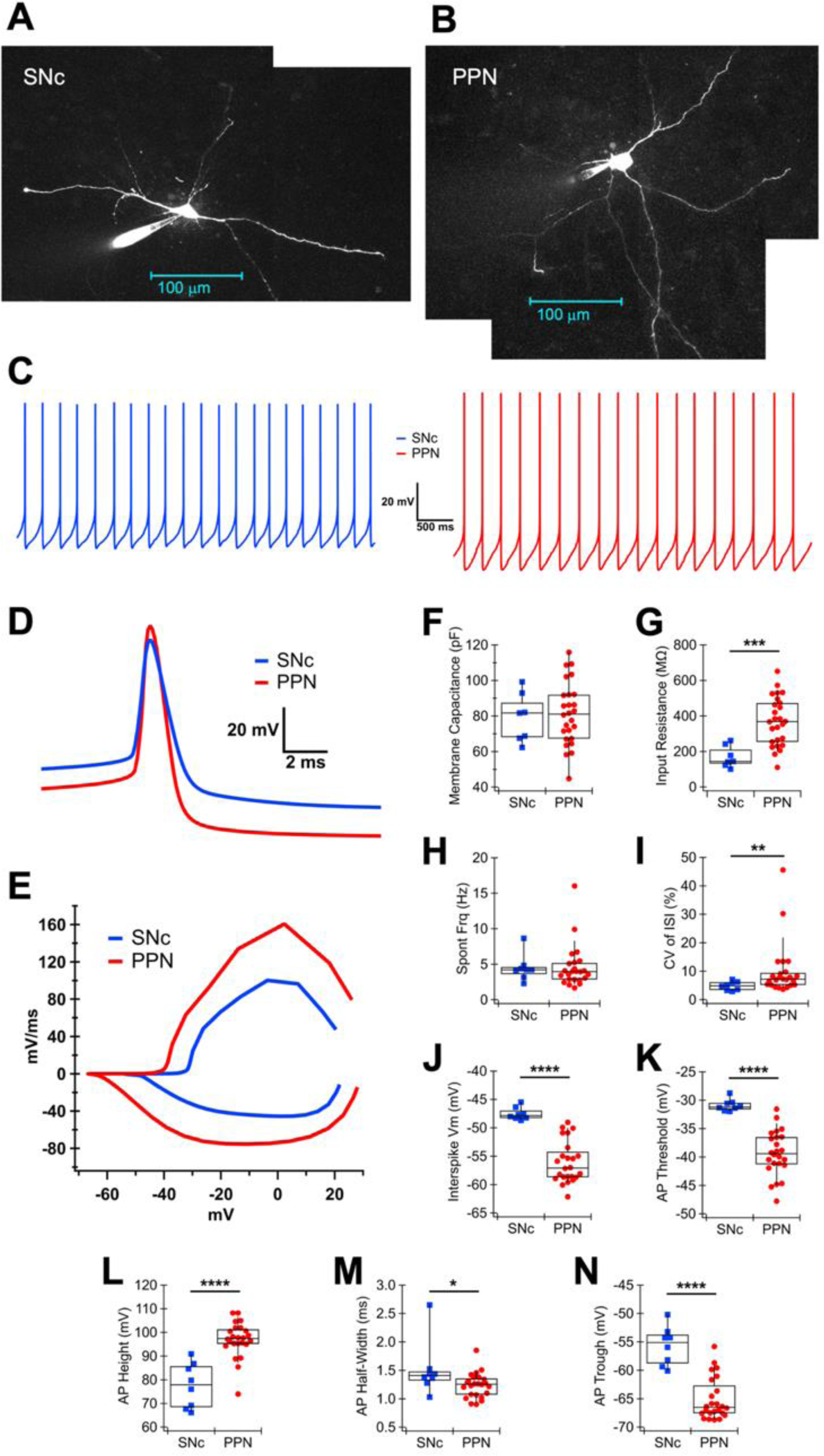
Electrophysiological characteristics of PPN and SNc neurons. **(A)** SNc dopaminergic neuron and **(B)** PPN cholinergic neuron with patch pipettes visualized from maximum intensity projection of Z-stacks using Alexa Fluor 594. **(C)** Representative whole-cell recordings of spontaneous pacemaking of a SNc dopaminergic neuron (blue) and a PPN cholinergic neuron (red). **(D)** Average action potential (AP) waveforms obtained from 30 seconds of pacemaking of the same cells in (C). **(E)** Phase plot of the average AP waveforms in (D). **(F)** Membrane capacitance and **(G)** input resistance of SNc (blue squares) and PPN (red circles) neurons measured using a −5 mV step from the holding potential of −70 mV. **(H)** Spontaneous firing frequency (no current injected) of SNc and PPN neurons. **(I)** The firing regularity, represented by the coefficient of variation (CV) of the interspike interval (ISI), **(J)** interspike membrane potential (mV), **(K)** AP threshold potential, **(L)** AP spike height, **(M)** AP half-width, and **(N)** AP trough (the lowest Vm during afterhyperpolarization) of spontaneously firing SNc and PPN neurons.

Both SNc dopaminergic (Grace and Onn, 1989; Johnson et al., 1992) and PPN cholinergic neurons (Takakusaki and Kitai, 1997) are spontaneously active in the *ex vivo* brain slices. In our preparation, all SNc neurons were actively firing in the cell-attached configuration. In contrast, most cholinergic PPN neurons were quiescent until the intracellular content was dialyzed ∼2 min after obtaining the whole-cell configuration. When regular pacemaking was stabilized, PPN neurons had a spontaneous firing rate of 4.58 ± 0.62 Hz (n = 24), which was comparable to the rate of SNc neurons at 4.48 ± 0.66 Hz (n = 8; Mann-Whitney-Wilcoxon test, p = 0.4284; Fig. 1C, H). While the pacemaking of both cell types appeared robust, the spontaneous firing of PPN neurons was less regular than that of SNc neurons (PPN CV of ISI: 9.97 ± 1.91%, n = 24; SNc CV of ISI: 4.87 ± 0.55%, n = 8; Mann-Whitney-Wilcoxon test, p = 0.0076; Fig. 1I).

Although SNc and PPN neurons did not differ in spontaneous firing rates, SNc neurons exhibited significantly more depolarized membrane potential (Vm) in the interspike interval than PPN neurons (SNc interspike Vm: −47.56 ± 0.39 mV, n = 8; PPN interspike Vm: −56.08 ± 0.75 mV, n = 24; Mann-Whitney-Wilcoxon test, p < 0.0001; Fig. 1J). When comparing the shape of action potentials (AP) (Fig. D, E), SNc neurons on average had a significantly more depolarized AP threshold (SNc AP threshold: - 30.93 ± 0.37 mV, n = 8; PPN AP threshold: −39.34 ± 0.80 mV, n = 24; Mann-Whitney-Wilcoxon test, p < 0.0001; Fig. 1K), a shorter spike height (SNc AP height: 77.65 ± 3.32 mV, n = 8; PPN AP height: 96.90 ± 1.51 mV, n = 24; Mann-Whitney-Wilcoxon test, p < 0.0001; Fig. 1L), a longer half-width (SNc AP half-width: 1.51 ± 0.17 ms, n = 8; PPN AP half-width: 1.24 ± 0.04 ms, n = 24; Mann-Whitney-Wilcoxon test, p = 0.0258; Fig. 1M), and a shallower afterhyperpolarization trough (SNc AP trough: −55.70 ± 1.19 mV, n = 8; PPN AP trough: −65.10 ± 0.75 mV, n = 24; Mann-Whitney-Wilcoxon test, p < 0.0001; Fig. 1N). Together, these observations of spontaneous firing showed that PPN cholinergic and SNc dopaminergic neurons engage in similar pacemaking activity; however, the differences in average Vm, firing regularity, and AP shape suggest that pacemaking may be governed by different ion channels in these two neuronal populations.

### Sodium channel blockage decreases tonic calcium in PPN cholinergic neurons

In SNc dopaminergic neurons, AP backpropagation into the dendrites can sensitively regulate Ca^2+^ entry (Hage and Khaliq, 2015; Wilson and Callaway, 2000) and neurotransmitter release (Beckstead et al., 2004; Gantz et al., 2013; Rice et al., 1997) in a frequency-dependent manner. Two major sources of Ca^2+^ account for the Ca^2+^ influx during firing activity: AP-evoked Ca^2+^ and Ca^2+^ entry at subthreshold voltages. Previous work has shown that subthreshold depolarization contributes greatly to dendritic Ca^2+^ levels even in the absence of firing in SNc dopaminergic neurons (Chan et al., 2007; Guzman et al., 2009; Hage and Khaliq, 2015). To test which mechanism mediates activity-associated Ca^2+^ increase in PPN cholinergic neurons and to directly compare the results with SNc dopaminergic neurons, we filled the patch pipette with the green Ca^2+^-sensitive dye Fluo-5F and imaged the soma and dendrites of PPN or SNc neurons during tonic and phasic firing. Linescans were taken at three sites on one cell (Fig. 2A): soma, proximal dendrite (≤50 μm), and distal dendrites (>50 μm). Ca^2+^ signals were calculated by dividing the changes in green fluorescence by red fluorescence and normalized to saturated Ca^2+^ conditions (presented as G/Gs, Fig. 2B). During tonic firing, a Ca^2+^ transient closely correlating to each peak of somatic AP (“pacemaking Ca^2+^”) could be observed in the soma of 3.8% (1 out of 26), in the proximal dendrites of 46% (12 out of 26), and in the distal dendrites of 42% (11 out of 26) of the imaged PPN neurons. The peaks of pacemaking Ca^2+^ were especially prominent in the dendrites of slow-firing cells (<3 Hz), but often undefinable in the soma or in faster-firing cells. To observe phasic Ca^2+^ entry, a 200-pA current step was applied to evoke burst-like firing and a robust Ca^2+^ transient during the linescan imaging (Fig. 1C).

**Figure 2.**
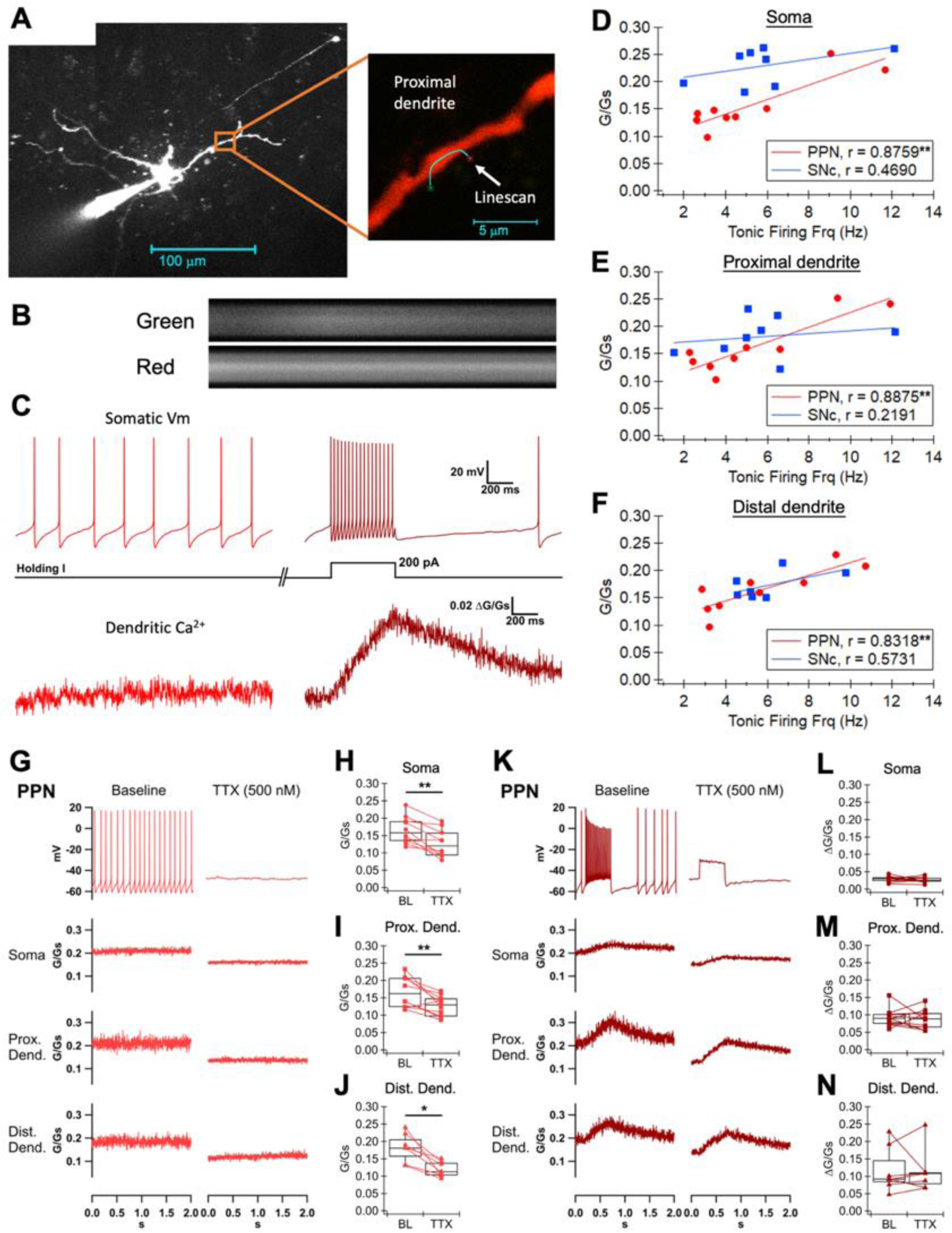
Dendritic calcium recordings in PPN cholinergic neurons with and without sodium channel blockade. **(A)** Representative image showing the morphology of a PPN cholinergic neuron with a patch pipette filled with Alexa Fluor 594 and Fluo-5F (left) and a zoomed-in image of the area indicated by the orange square showing the site of linescan taken at the proximal dendrite of the neuron (right). **(B)** Linescan fluorescence signals (separated into green and red channels) showing a Ca^2+^ transient during a 200-pA current step measured at the dendrite in (A). **(C)** Time-matched whole-cell somatic Vm recording and dendritic linescan Ca^2+^ signal during 0 pA holding current (light red) and a 200-pA current step (dark red), measured at the dendrite in (A). In this representative recording, pacemaking Ca^2+^ was observed in the basal condition (0 pA). **(D)** The basal Ca^2+^ level measured at the soma, **(E)** proximal dendrite (≤50 μm), and **(F)** distal dendrite (>50 μm) plotted against the tonic firing frequency measured at the soma of PPN (red circles) and SNc (blue squares) neurons. The data from one cell type at each location were fitted to linear regression. The corresponding Pearson correlation coefficients (r) and statistical significance are shown. **(G)** Representative time-matched recordings of somatic Vm and Ca^2+^ signals measured at the soma, proximal dendrite, and distal dendrite of a PPN cholinergic neuron during 0 pA holding current at the baseline and after bath treatment of tetrodotoxin (TTX, 500 nM). **(H)** Summary box plot of the basal Ca^2+^ levels measured at the soma, **(I)** proximal dendrite, and **(J)** distal dendrite at the baseline (BL) and after TTX treatment. **(K)** Representative time-matched recordings of somatic Vm and Ca^2+^ signals measured at the soma, proximal dendrite, and distal dendrite of a PPN cholinergic neuron during 200-pA current step at the baseline and after bath treatment of TTX. **(L)** Summary box plot of the phasic Ca^2+^ amplitudes (peak − basal Ca^2+^) measured at the soma, **(M)** proximal dendrite, and **(N)** distal dendrite at the baseline and after TTX treatment.

To evaluate the relationship between somatodendritic Ca^2+^ and AP frequency in PPN neurons, we plotted the basal Ca^2+^ levels against the simultaneous tonic firing frequency of PPN neurons alongside the SNc neuron data for comparison. The data from one cell type at each cellular compartment were fitted to linear regression. Interestingly, basal Ca^2+^ levels in PPN neurons were highly correlated with firing rates in all somatodendritic compartments, having a statistically significant correlation coefficient (r) value of 0.8759 at the soma (n = 9, p = 0.0020; Fig. 2D), 0.8875 at the proximal dendrites (n = 9, p = 0.0014; Fig. 2E), and 0.8318 at the distal dendrites (n = 9, p = 0.0054; Fig 2F). The relationship between basal Ca^2+^ levels and firing rates showed less correlation and did not reach statistical significance in all somatodendritic compartments of SNc neurons. The r value at the SNc soma was 0.4690 (n = 8, p = 0.2411), 0.2191 at the proximal dendrite (n = 8, p = 0.6022), and 0.5731 at the distal dendrite (n = 7, p = 0.1787). The lower r values in the SNc are consistent with previous findings that dendritic Ca^2+^ in SNc dopaminergic neurons are mostly induced by subthreshold depolarization and therefore not necessarily correlated with AP firing, whereas the high r values in the PPN suggest most of the Ca^2+^ is AP-evoked and frequency-dependent.

We tested the hypothesis that dendritic Ca^2+^ in the PPN is AP-dependent by silencing the sodium channel-mediated spikes with TTX. After bath application of TTX (500 nM), AP spiking in PPN neurons was effectively silenced, revealing a stable and rather depolarized Vm (Fig. 2G). This inhibition of tonic firing significantly decreased the basal Ca^2+^ levels in the soma from 0.165 ± 0.012 to 0.127 ± 0.012 G/Gs (n = 10, Wilcoxon signed-rank test, p = 0.0020; Fig. 2H), proximal dendrites from 0.168 ± 0.014 to 0.127 ± 0.009 G/Gs (n = 10, Wilcoxon signed-rank test, p = 0.0059; Fig. 2I), and distal dendrites from 0.182 ± 0.016 to 0.120 ± 0.009 G/Gs (n = 7, Wilcoxon signed-rank test, p = 0.0488; Fig. 2J). In contrast, during evoked phasic firing, TTX caused a drop in basal Ca^2+^ levels but the amplitudes of depolarization-induced Ca^2+^ transients (peak − basal Ca^2+^) were not changed in any somatodendritic compartment (Wilcoxon signed-rank test, phasic Ca^2+^ before vs. after TTX; soma: n = 10, p = 0.5566; proximal dendrite: n = 10, p = 0.6953; distal dendrite: n = 7, p = 0.8125; Fig. 2K-N). These results suggest that, similar to SNc dopaminergic neurons, PPN cholinergic neurons exhibit pacemaking Ca^2+^ that corresponds to somatic APs. AP firing contributes to a significant component of somatodendritic Ca^2+^ during tonic firing in PPN neurons, but the Ca^2+^ entry during phasic depolarization does not require AP firing or the activation of TTX-sensitive sodium channels.

### L-type calcium channel blockage has minimal effects on action potential kinetics in PPN cholinergic neurons

Because TTX inhibition of AP firing in PPN neurons caused a significant reduction in basal Ca^2+^ levels, we hypothesized that most of the tonic activity-associated Ca^2+^ influx occurs during the AP. This is in contrast to SNc neurons, in which the low-threshold L-type Ca^2+^ channel Cav1.3 activates at subthreshold voltages and mediates most of the Ca^2+^ influx during pacemaking (Chan et al., 2007; Philippart et al., 2016; Puopolo et al., 2007). To further investigate the mechanisms mediating activity-associated Ca^2+^ entry in PPN neurons, and whether L-type Ca^2+^ channels play a similar role in the regulation of AP kinetics in SNc and PPN neurons, we treated the two cell types with the L-type Ca^2+^ channel blocker nifedipine and compared the changes in their AP properties.

Consistent with pacemaking Ca^2+^ in SNc neurons depending on L-type Ca^2+^ channels, nifedipine (10 μM) bath treatment caused a modest but significant reduction in tonic firing rate from 5.74 ± 0.94 to 4.60 ± 0.74 Hz in SNc neurons (n = 8, Wilcoxon signed-rank test, p = 0.039; Fig. 3C). Accompanying the decrease in firing rate, there was a widening of the AP spike, from 1.70 ± 0.08 to 2.09 ± 0.16 ms in the half-width (n = 8, Wilcoxon signed-rank test, p = 0.023; Fig. 3D); this slowing of AP kinetics is clearly visible in the representative AP waveform (Fig. 3A) and phase plot (Fig. 3B). However, there were no significant changes in the interspike Vm (n = 8, Wilcoxon signed-rank test, p = 0.2500; Fig. 3E) and the depth of afterhyperpolarization trough (n = 8, Wilcoxon signed-rank test, p = 0.148; Fig. 3F). The firing regularity (n = 8, Wilcoxon signed-rank test, p = 0.2500), AP threshold (n = 8, Wilcoxon signed-rank test, p = 0.7422), and spike height (n = 8, Wilcoxon signed-rank test, p = 0.1094) were also not affected by the nifedipine treatment (Supplementary Fig. S1).

**Figure 3.**
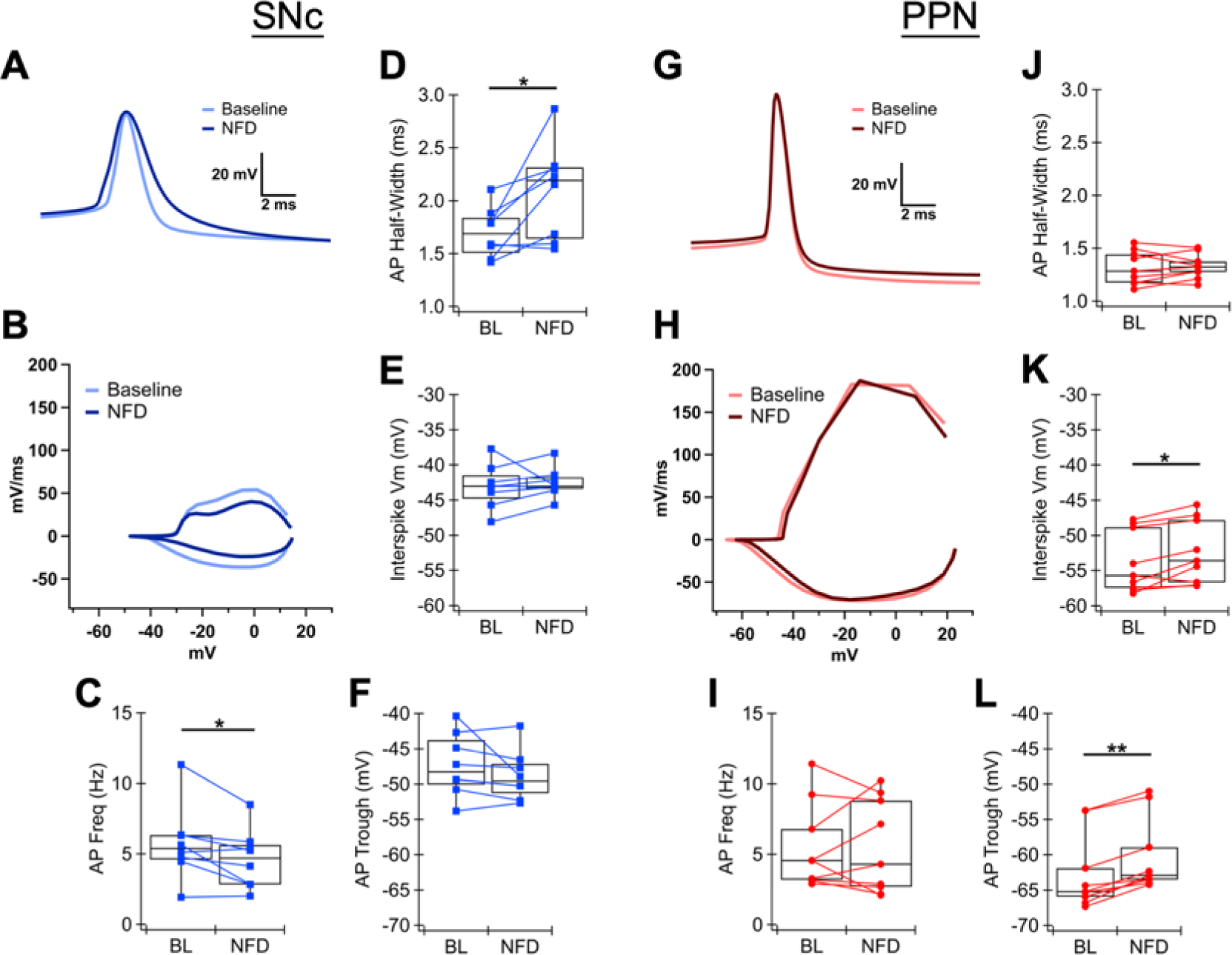
L-type calcium channel regulation of action potential shape in PPN and SNc neurons. **(A)** Average AP waveform of a SNc dopaminergic neuron obtained from 30-s of tonic firing at the baseline (BL, light blue) and after bath treatment of nifedipine (NFD, 10 μM; dark blue). **(B)** Phase plot of the average AP waveforms in (A). **(C)** Tonic firing frequency, **(D)** AP half-width, **(E)** interspike Vm, and **(F)** afterhyperpolarization trough of SNc neurons at the baseline and after nifedipine treatment. **(G)** Average AP waveform of a PPN cholinergic neuron obtained from 30-s of tonic firing at the baseline (light red) and after bath treatment of nifedipine (dark red). **(H)** Phase plot of the average AP waveforms in (G). **(I)** Tonic firing frequency, **(J)** AP half-width, **(K)** interspike Vm, and **(L)** afterhyperpolarization trough of SNc neurons at the baseline and after nifedipine treatment.

However, unlike the SNc neurons, nifedipine did not affect the tonic firing frequency (n = 9, Wilcoxon signed-rank test, p = 1.000; Fig. 3I) or AP half-width (n = 9, Wilcoxon signed-rank test, p = 0.8203; Fig. 3J) of PPN neurons. As shown by the representative AP waveform (Fig. 3G) and phase plot (Fig. 3H), there were minimal changes to the overall AP shape, except a mild but significant depolarization of the interspike Vm from −53.84 ± 1.46 to −52.38 ± 1.50 mV (n = 9, Wilcoxon signed-rank test, p = 0.0195; Fig. 3K) and reduction of afterhyperpolarization trough from −62.69 ± 1.77 to −60.23 ± 1.75 mV (n = 9, Wilcoxon signed-rank test, p = 0.0004; Fig. 3L). The firing regularity (n = 9, Wilcoxon signed-rank test, p = 0.0742), AP threshold (n = 9, Wilcoxon signed-rank test, p = 0.2031), and spike height (n = 9, Wilcoxon signed-rank test, p = 0.0547) were unaffected by the nifedipine treatment (Supplementary Fig. S1). These results show that the effects of L-type Ca^2+^ channel blockage are distinct in those two cell types: nifedipine slows the kinetics of the AP spike in SNc neurons, whereas in the PPN neurons, nifedipine depolarizes the interspike Vm.

### L-type calcium channel blockage does not reduce tonic calcium in PPN cholinergic neurons

We established that blockage of L-type Ca^2+^ channels affected the kinetics of tonic firing to a lesser extent in PPN neurons than in SNc neurons. This suggests that the pacemaking of PPN neurons may be less reliant on L-type Ca^2+^ conductance, and L-type Ca^2+^ current likely accounts for a smaller portion of depolarization-induced Ca^2+^ influx in PPN neurons. To determine whether L-type Ca^2+^ channels play a significant role in pacemaking Ca^2+^ in PPN neurons, we took two-photon dendritic linescan imaging of tonically firing PPN neurons before and after nifedipine treatment, and compared the change in tonic Ca^2+^ levels to that of SNc neurons.

PPN neurons exhibited heterogeneous responses to nifedipine bath treatment. In one PPN neuron, whose tonic firing rate (from 3.67 to 3.81 Hz) and firing regularity (CV of ISI from 43.23 to 45.07%) were almost identical before and after nifedipine, the basal Ca^2+^ level in the proximal dendrite also remained constant (from 0.127 to 0.128 G/Gs; Fig. 4A). The dendritic Ca^2+^ oscillated with AP spiking, but the peaks were less defined and only slightly reduced in nifedipine. In another PPN neuron, the tonic firing became much slower (from 4.69 to 1.66 Hz) and more irregular (CV of ISI from 8.30 to 25.30%) after nifedipine treatment, and the basal Ca^2+^ level in the proximal dendrite was likewise reduced (from 0.141 to 0.120 G/Gs; Fig. 4B). This cell exhibited clear pacemaking Ca^2+^, having well-defined peaks that neatly correlated with AP spiking and were greatly reduced by nifedipine treatment. Therefore, we observed that some PPN cholinergic neurons appear sensitive to nifedipine, while others are unresponsive. When the results were averaged across all PPN cells, there were no significant changes in the basal Ca^2+^ levels after nifedipine in any somatodendritic compartment during tonic firing (Wilcoxon signed-rank test, tonic Ca^2+^ before vs. after nifedipine; soma: n = 10, p = 0.1934; proximal dendrite: n = 10, p = 0.1309; distal dendrite: n = 10, p = 0.1309; Fig. 4C-F). The unaltered tonic Ca^2+^ levels correspond to the lack of change in the averaged firing frequency in PPN neurons after L-type Ca^2+^ channel blockage.

**Figure 4.**
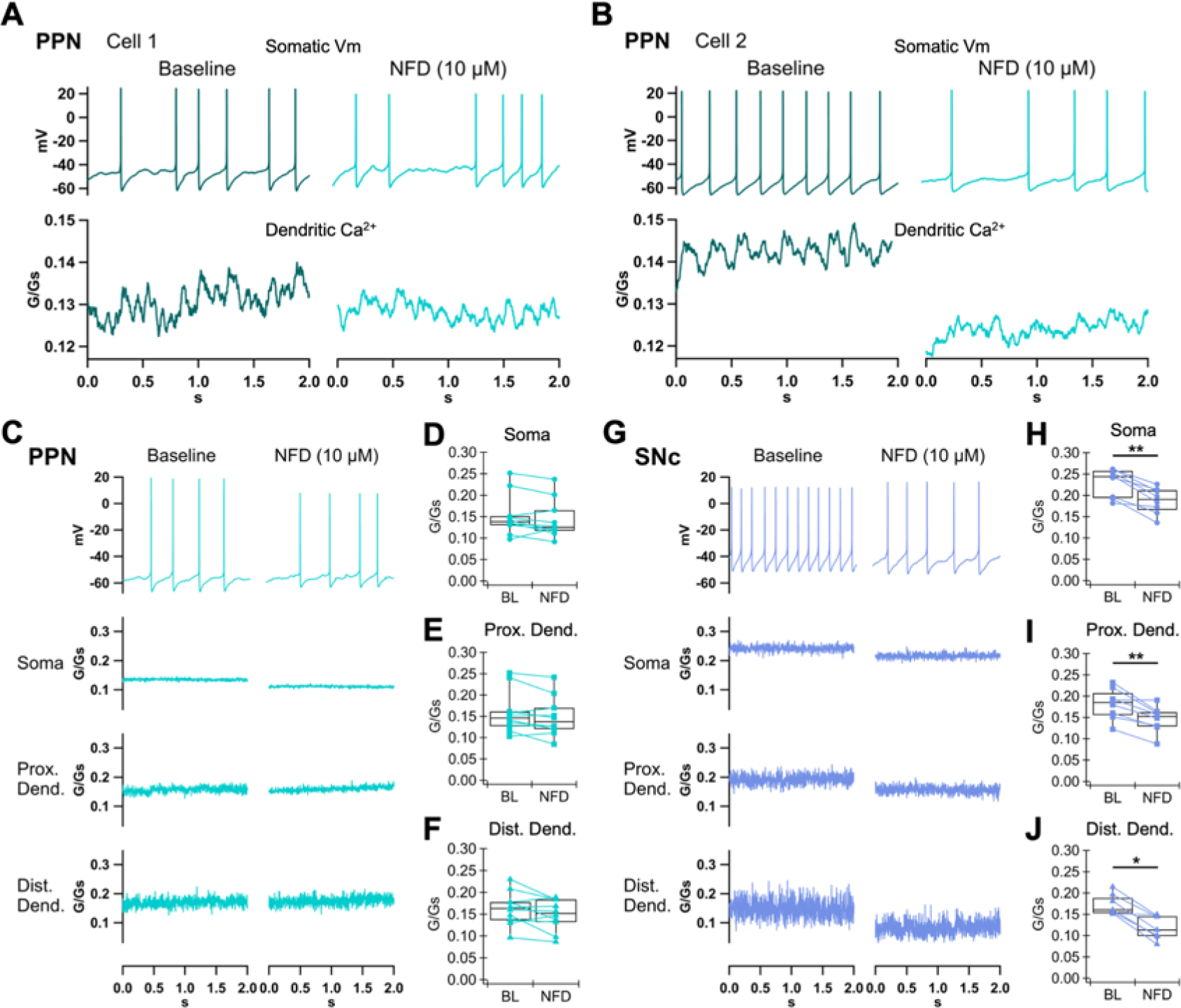
Contribution of L-type channels to PPN and SNc tonic calcium. **(A)** Example PPN cholinergic neuron whose tonic firing rate and basal Ca^2+^ level measured at the proximal dendrite did not change after nifedipine (NFD, 10 μM) bath treatment. **(B)** Example PPN cholinergic neuron whose tonic firing rate and basal Ca^2+^ level measured at the proximal dendrite both decreased after nifedipine treatment. The Ca^2+^ traces in (A) and (B) were smoothed using the boxcar method with a factor of 20 points. Pacemaking Ca^2+^ that correlated with AP spiking could be seen in the baseline condition. **(C)** Representative time-matched recordings of somatic Vm and Ca^2+^ signals measured at the soma, proximal dendrite, and distal dendrite of a PPN cholinergic neuron during 0 pA holding current at the baseline and after nifedipine treatment. **(D)** Summary box plot of the basal Ca^2+^ levels measured at the soma, **(E)** proximal dendrite, and **(F)** distal dendrite of PPN cholinergic neurons at the baseline (BL) and after nifedipine treatment. **(G)** Representative time-matched recordings of somatic Vm and Ca^2+^ signals measured at the soma, proximal dendrite, and distal dendrite of a SNc dopaminergic neuron during 0 pA holding current at the baseline and after nifedipine treatment. **(H)** Summary box plot of the basal Ca^2+^ levels measured at the soma, **(I)** proximal dendrite, and **(J)** distal dendrite of SNc dopaminergic neurons at the baseline and after nifedipine treatment.

Because there is a well-defined role for low-threshold L-type Ca^2+^ channels (Cav1.3) in SNc dopaminergic neuron tonic Ca^2+^ levels (Chan et al., 2007; Guzman et al., 2009; Hage and Khaliq, 2015; Puopolo et al., 2007), we ran the same experiments on SNc neurons. In contrast to the lack of effect in PPN neurons, nifedipine exerted a significant and consistent effect on the tonic Ca^2+^ levels in SNc neurons (Fig. 4G). Accompanying the slowing of tonic firing frequency and AP kinetics, the basal Ca^2+^ levels in the soma (from 0.229 ± 0.012 to 0.187 ± 0.011 G/Gs, n = 8, Wilcoxon signed-rank test, p = 0.0078; Fig. 4H), proximal dendrites (from 0.181 ± 0.013 to 0.146 ± 0.011 G/Gs, n = 8, Wilcoxon signed-rank test, p = 0.0078; Fig. 4I), and distal dendrites (from 0.173 ± 0.009 to 0.119 ± 0.011 G/Gs, n = 7, Wilcoxon signed-rank test, p = 0.0156; Fig. 4J) of SNc neurons were all significantly decreased by nifedipine. This trend was strong and consistent across cells, as all 8 SNc neurons recorded showed decreases in tonic Ca^2+^ in all somatodendritic compartments after nifedipine treatment. Therefore, in the same recording conditions, L-type Ca^2+^ channel blockage reduces tonic dendritic Ca^2+^ in SNc dopaminergic neurons, but not in PPN cholinergic neurons.

### L-type calcium channel blockage decreases phasic calcium in PPN cholinergic neurons

To determine whether L-type Ca^2+^ channels play a significant role in depolarization-induced Ca^2+^ entry during phasic firing in PPN neurons, we applied a 200-pA current step to elicit a burst of fast phasic firing and elevation of somatodendritic Ca^2+^. Nifedipine significantly decreased the amplitudes of phasic firing-induced Ca^2+^ entry in all somatodendritic compartments in the PPN neurons (Fig. 5A), from 0.033 ± 0.003 to 0.028 ± 0.003 ΔG/Gs in the soma (n = 10, Wilcoxon signed-rank test, p = 0.0195; Fig. 5B), 0.104 ± 0.011 to 0.081 ± 0.012 ΔG/Gs in the proximal dendrites (n = 10, Wilcoxon signed-rank test, p = 0.0137; Fig. 5C), and 0.109 ± 0.010 to 0.072 ± 0.008 ΔG/Gs in the distal dendrites (n = 10, Wilcoxon signed-rank test, p = 0.0020; Fig. 5D). DMSO (0.05%) alone had no effects on phasic Ca^2+^ in the soma (n = 6, Wilcoxon signed-rank test, p = 0.1563) and proximal dendrites (n = 6, Wilcoxon signed-rank test, p = 0.5625) of PPN neurons, while there was a statistically significant decrease of phasic Ca^2+^ amplitudes in the distal dendrite (n = 6, Wilcoxon signed-rank test, p = 0.0313; Supplementary Fig. S2). This indicates that the phasic Ca^2+^ signal in the distal dendrites undergoes rundown over time. The efficacy of nifedipine in reducing phasic Ca^2+^ in the soma and proximal dendrites suggests that fast burst-like firing in PPN neurons can reliably activate L-type Ca^2+^ channels, and that these channels are likely to be the high-threshold (Cav1.2) subtype.

**Figure 5.**
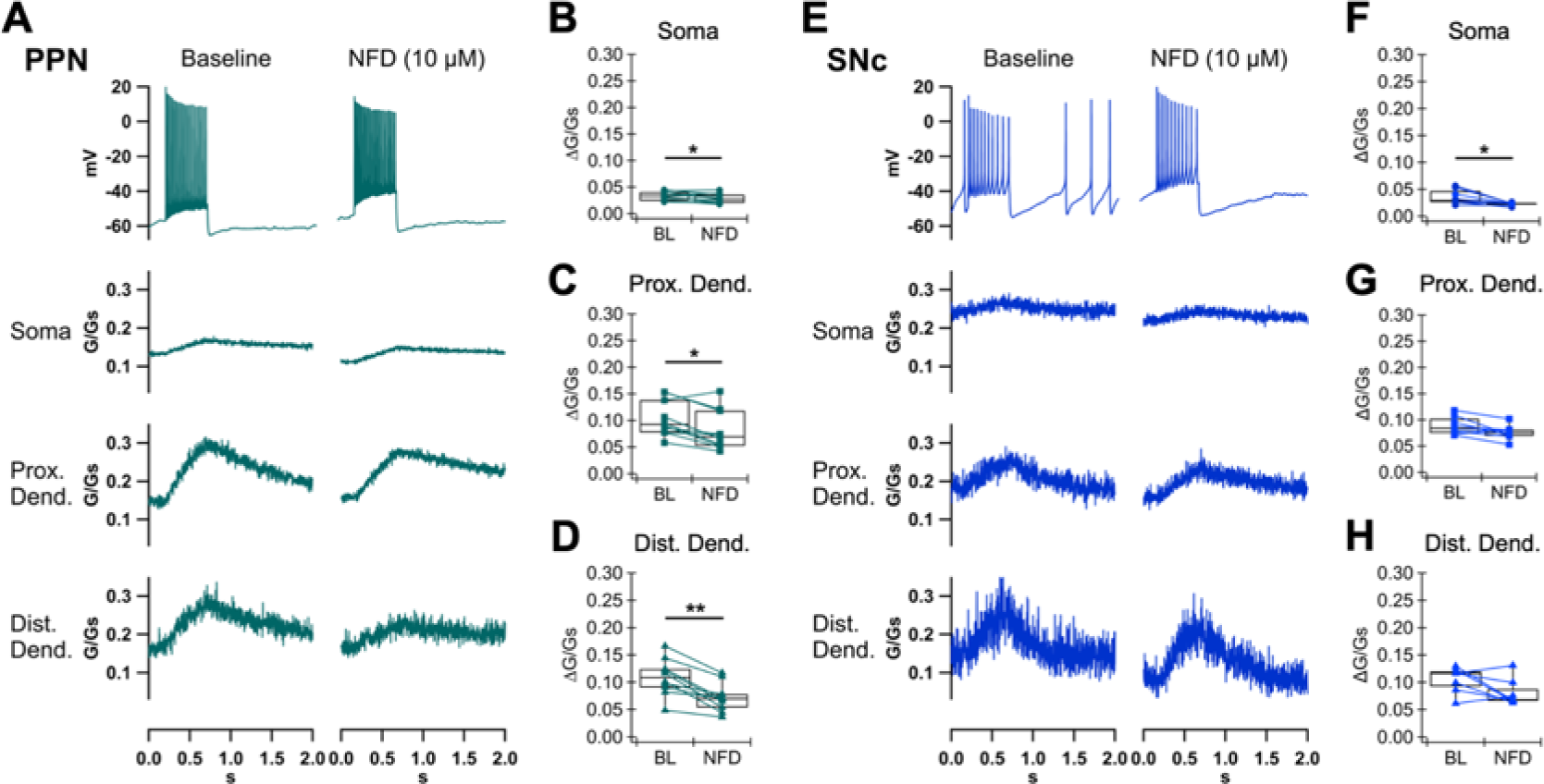
Contribution of L-type channels to PPN and SNc phasic calcium. **(A)** Representative time-matched recordings of somatic Vm and Ca^2+^ signals measured at the soma, proximal dendrite, and distal dendrite of a PPN cholinergic neuron during 200-pA current step at the baseline and after bath treatment of nifedipine (NFD). **(B)** Summary box plot of the phasic Ca^2+^ amplitudes (peak − basal Ca^2+^) measured at the soma, **(C)** proximal dendrite, and **(D)** distal dendrite of PPN cholinergic neurons at the baseline (BL) and after nifedipine treatment. **(E)** Representative time-matched recordings of somatic Vm and Ca^2+^ signals measured at the soma, proximal dendrite, and distal dendrite of a SNc dopaminergic neuron during 200-pA current step at the baseline and after bath treatment of nifedipine. **(F)** Summary box plot of the phasic Ca^2+^ amplitudes measured at the soma, **(G)** proximal dendrite, and **(H)** distal dendrite of SNc dopaminergic neurons at the baseline and after nifedipine treatment.

When the same protocol was performed in SNc neurons (Fig. 5E), we found that nifedipine significantly reduced the amplitude of phasic Ca^2+^ in the soma, from 0.035 ± 0.005 to 0.023 ± 0.001 ΔG/Gs (n = 8, Wilcoxon signed-rank test, p = 0.0156; Fig. 5F). Phasic Ca^2+^ in the proximal and distal dendrites showed trends of decrease in nifedipine, but the effects did not reach statistical significance (proximal dendrite: n = 8, Wilcoxon signed-rank test, p = 0.0547; distal dendrites: n = 7, Wilcoxon signed-rank test, p = 0.0781; Fig. 5G-H). This result supports the idea that Cav1.2 is more weakly expressed than the low-threshold Cav1.3 L-type channels in SNc dopaminergic neurons (Chan et al., 2007; Dufour et al., 2014; Philippart et al., 2016). Together, our results suggest that L-type Ca^2+^ channels play opposing roles in tonic vs. phasic Ca^2+^ entry in PPN and SNc neurons: while L-type Ca^2+^ channels contribute significantly to tonic Ca^2+^ but less to phasic Ca^2+^ in SNc neurons, the same family of channels account for a significant amount of phasic Ca^2+^ but not tonic Ca^2+^ in PPN neurons.

### L-type calcium channels contribute to tonic calcium levels throughout the dendrites in SNc but not PPN neurons

Previous studies showed that depolarization-induced Ca^2+^ signals, which could be evoked by AP backpropagation, travel along the dendrites of SNc dopaminergic neurons and decay very little with distance (Hage and Khaliq, 2015). This is due to strong electronic coupling of the soma and the dendrites, as well as the presence of Ca^2+^ channels active at subthreshold potentials. In this study, we found that PPN neurons have higher input resistance than SNc neurons, suggesting even tighter electronic coupling throughout the cell. However, PPN neurons appear to express only high-threshold L-type Ca^2+^ channels, which would require significant depolarization invading into the dendrites to activate. To determine the amounts of L-type Ca^2+^ channel-mediated Ca^2+^ influx throughout the dendrites, we evaluated the nifedipine-dependent reduction in tonic Ca^2+^ signals between SNc and PPN neurons across distances from the soma.

To directly compare the role of L-type Ca^2+^ channels in SNc and PPN dendrites, we grouped the Ca^2+^ data into three compartments: soma, proximal dendrite, and distal dendrite (Fig. 6G). In the box plots, the ΔG/Gs value is calculated from subtracting the Ca^2+^ level after nifedipine treatment from the control Ca^2+^ level (Ca^2+^ before nifedipine − after nifedipine); therefore, a positive ΔG/Gs indicates a decrease of Ca^2+^ level by nifedipine while a negative ΔG/Gs indicates an increase. The reduction of basal Ca^2+^ during tonic firing due to nifedipine was significantly larger in the soma of SNc neurons than in the soma of PPN neurons (SNc soma: 0.042 ± 0.006 ΔG/Gs, n = 8; PPN soma: 0.009 ± 0.005 ΔG/Gs, n = 10; Mann-Whitney-Wilcoxon test, p = 0.0021; Fig. 6A). Similarly, nifedipine caused a significantly larger reduction of tonic Ca^2+^ in the proximal and distal dendrites of SNc neurons, compared to the proximal and distal dendrites of PPN neurons, respectively (SNc proximal dendrites: 0.036 ± 0.008 ΔG/Gs, n = 8; PPN proximal dendrites: 0.010 ± 0.005 ΔG/Gs, n = 10; Mann-Whitney-Wilcoxon test, p = 0.0117; Fig. 6B. SNc distal dendrites: 0.054 ± 0.009 ΔG/Gs, n = 7; PPN distal dendrites: 0.015 ± 0.008 ΔG/Gs, n = 10; Mann-Whitney-Wilcoxon test, p = 0.0046; Fig. 6C). By contrast, when the reduction of phasic firing-evoked Ca^2+^ due to nifedipine was compared, there was no significant difference between SNc and PPN neurons at the soma (SNc: n = 8; PPN: n = 10; Mann-Whitney-Wilcoxon test, p = 0.3599; Fig. 6D), proximal dendrites (SNc: n = 8; PPN: n = 10; Mann-Whitney-Wilcoxon test, p = 0.2031; Fig. 6E), or distal dendrites (SNc: n = 7; PPN: n = 10; Mann-Whitney-Wilcoxon test, p = 0.4747; Fig. 6F). Thus, our results show that nifedipine has consistently larger effects on tonic Ca^2+^ in SNc neurons regardless of distances from the soma. This suggests that SNc neurons have larger L-type Ca^2+^ channel-mediated tonic influx throughout the dendrites compared to PPN neurons.

**Figure 6.**
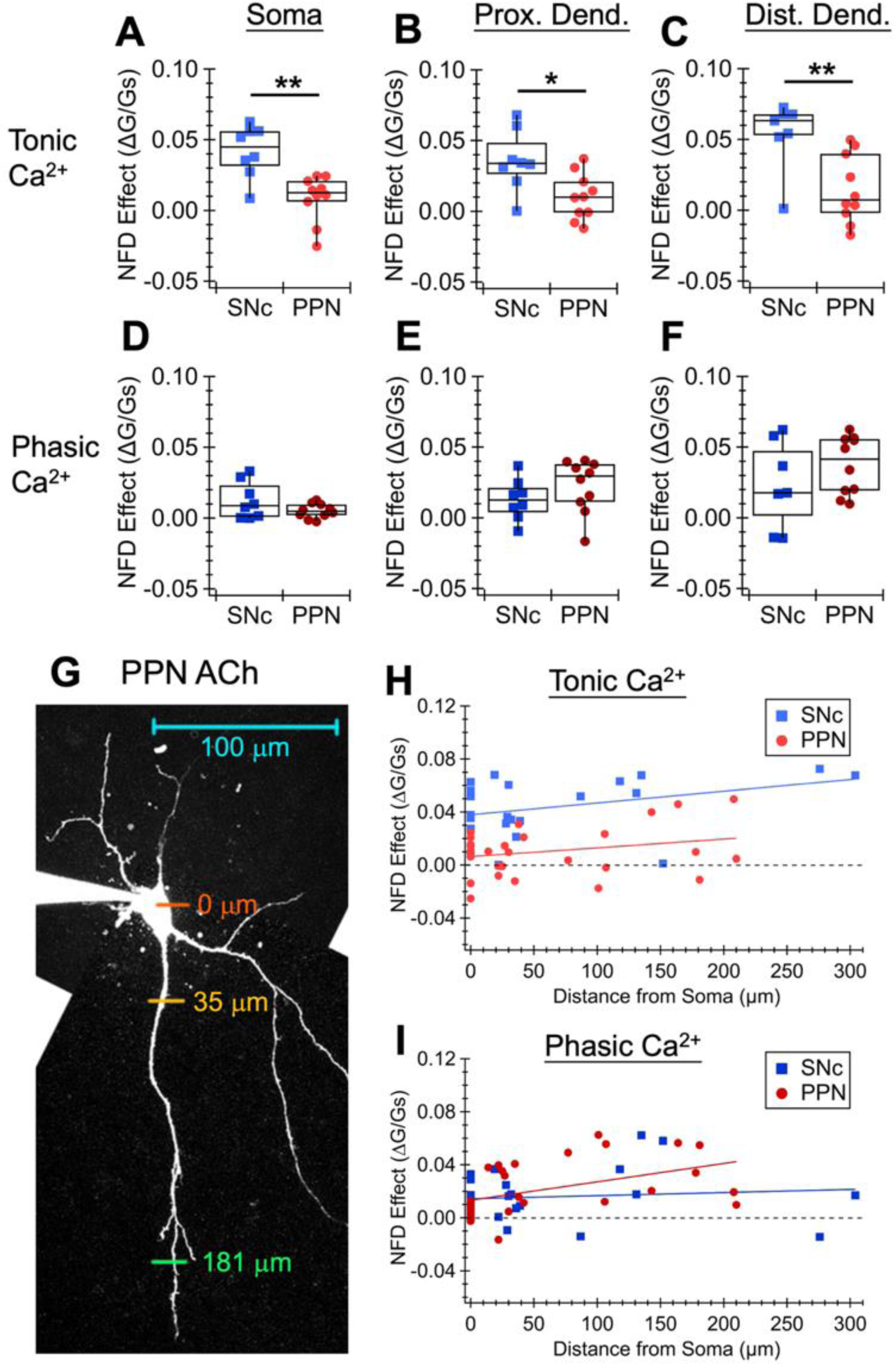
Contribution of L-type channels throughout the dendrites of PPN and SNc neurons. **(A)** The difference in the basal Ca^2+^ levels (during tonic firing at 0 pA holding current) before and after bath treatment of nifedipine (Ca^2+^ before NFD − after NFD) in SNc dopaminergic (light blue squares) and PPN cholinergic (light red circles) neurons, measured at the soma, **(B)** proximal dendrite, and **(C)** distal dendrite. **(D)** The difference in the amplitudes of phasic Ca^2+^ evoked by a 200-pA current step before and after bath treatment of nifedipine in SNc dopaminergic (dark blue squares) and PPN cholinergic (dark red circles) neurons, measured at the soma, **(E)** proximal dendrite, and **(F)** distal dendrite. **(G)** Example linescan sites, indicated by colored bars, taken at the soma (orange), proximal dendrite (yellow), and distal dendrite (green) of a PPN cholinergic neuron. **(H)** The difference in the basal Ca^2+^ levels before and after nifedipine treatment in SNc dopaminergic (light blue squares) and PPN cholinergic (light red circles) neurons plotted against distance from the soma of the linescan site. **(I)** The difference in the amplitudes of phasic Ca^2+^ before and after nifedipine treatment in SNc dopaminergic (dark blue squares) and PPN cholinergic (dark red circles) neurons plotted against distance from the soma of the linescan site. The data from each cell type were fitted to linear regression.

To more precisely evaluate the role of L-type Ca^2+^ channels across the extent of the dendritic arbor, we plotted the Ca^2+^ reduction due to nifedipine by the distances from the soma. We found that the tonic Ca^2+^ reduction due to nifedipine was consistently larger in SNc neurons compared to PPN neurons throughout the extent of the dendritic arbor. This is demonstrated by the SNc trendline having a much larger y-intercept value of 0.038 ± 0.001 ΔG/Gs than that of the PPN trendline, 0.006 ± 0.001 ΔG/Gs, while the two trendlines have similar slopes: 8.86 × 10^−5^ ± 1.06 × 10^−5^ (ΔG/Gs)/µm for SNc and 6.65 × 10^−5^ ± 0.88 × 10^−5^ (ΔG/Gs)/µm for PPN (Fig. 6H). These data indicate that L-type Ca^2+^channels (likely the low-threshold Cav1.3 subtype) contribute to tonic pacemaking Ca^2+^ influx throughout the somatodendritic extent of SNc neurons, but do not contribute to tonic Ca^2+^ influx in any compartment of the PPN neurons.

Evaluating the nifedipine effect on phasic Ca^2+^, we found no difference in SNc and PPN neurons at locations closer to the soma, shown by the SNc and PPN trendlines crossing near the y-intercepts (SNc y-intercept: 0.014 ± 0.001 ΔG/Gs; PPN y-intercept: 0.013 ± 0.001 ΔG/Gs; Fig. 6I). In the distal dendrites, the PPN trendline has a steeper slope value of 1.38 × 10^−4^ ± 0.09 × 10^−4^ (ΔG/Gs)/µm, which was almost 10 times of the slope value of the SNc trendline, 2.32 × 10^−5^ ± 1.05 × 10^−5^ (ΔG/Gs)/µm. These results indicate that L-type Ca^2+^ channels (likely the high-threshold Cav1.2 subtype) contribute to phasic Ca^2+^ influx more substantially in the distal dendrites of PPN neurons, whereas these high-threshold channels make minimal but uniform contributions to phasic Ca^2+^ influx across SNc somatodendritic compartments.Together, our findings show that L-type Ca^2+^ channels contribute to phasic, but not tonic Ca^2+^ levels in PPN cholinergic neurons.

## Discussion

Our study finds that PPN cholinergic neurons display similar spontaneous firing properties and pacemaking-induced Ca^2+^ oscillations as SNc dopaminergic neurons, but that, unlike SNc neurons, tonic Ca^2+^ entry in PPN neurons is not mediated by L-type Ca^2+^ conductance. Since somatodendritic tonic Ca^2+^ levels in PPN neurons are strongly decreased by TTX-induced sodium channel blockage, but not by nifedipine-induced L-type Ca^2+^ channel blockage, our data indicate that most of the tonic Ca^2+^ in PPN neurons is AP-evoked. In contrast, burst-like phasic firing-induced Ca^2+^ transients in PPN neurons were significantly suppressed by nifedipine, supporting the idea that PPN neurons selectively express high-threshold L-type Ca^2+^ channels. In addition, we show that L-type channel blockage slowed the kinetics of APs in SNc dopaminergic neurons but had minimal effects on PPN cholinergic neurons, indicating the pacemaking activities of these two midbrain neuronal populations are regulated by fundamentally different ionic mechanisms.

The majority of research investigating cellular mechanisms in Parkinson’s disease has been done on SNc dopaminergic neurons, while there is a lack of understanding of the relevant basic physiological properties of PPN cholinergic neurons. Takakusaki & Kitai were the first to report high-threshold somatic Ca^2+^ oscillations mediated by L- and N-type Ca^2+^ channels in the PPN cholinergic neurons of male adolescent rats (Takakusaki and Kitai, 1997). More than a decade later, work from the Garcia-Rill Lab identified N- and P/Q-type Ca^2+^ channels as the ionic mechanisms underlying high-threshold somatic Ca^2+^ oscillations in PPN cholinergic neurons in young rats (Kezunovic et al., 2011), and imaged these high-threshold Ca^2+^ oscillations using ratiometric fluorescence Ca^2+^ sensors (Hyde et al., 2013). Although they showed that a depolarizing current ramp evoked Ca^2+^ transient in the soma as well as the proximal dendrites, suggesting the presence of Ca^2+^ channels throughout the cell, they did not measure Ca^2+^ in distal dendrites. Here, we measured activity-associated Ca^2+^ entry in distal dendrites up to 300 μm away from the soma, and directly compared PPN and SNc neurons in the same experimental design. We found that tonic Ca^2+^ in PPN neurons is depended on sodium channel-mediated spiking, whereas the propagation of phasic Ca^2+^ into the dendrites does not require AP firing. This is consistent with previous findings that the somatic Ca^2+^ transients induced by a depolarizing current ramp were unchanged or even larger in the presence of TTX (Hyde et al., 2013). We also found that nifedipine did not reduce tonic Ca^2+^ but inhibited a significant portion of phasic Ca^2+^ in PPN neurons. In the perspective of past findings, intracellular Ca^2+^ levels of PPN neurons during tonic firing and the nifedipine-insensitive portion of phasic Ca^2+^ likely depend on other high-threshold Ca^2+^ channels, namely the N- and P/Q-types.

In addition to imaging dendritic Ca^2+^, we investigate the effects of L-type channel blockage on PPN neuron AP kinetics. Our results show that the only significant effect of nifedipine on PPN neuron AP shape was a slight depolarization of the interspike Vm, including the afterhyperpolarization trough. Surprisingly, while nifedipine significantly decreased firing frequency and increased spike width in SNc neurons, there was no change in the interspike Vm. We suggest that these distinct nifedipine effects on the membrane potential could be caused by the interplay of different L-type Ca^2+^ channel subtypes and Ca^2+^-activated potassium channels. In both PPN and SNc neurons, apamin-sensitive SK channels have been reported to underlie the afterhyperpolarization phase of AP or Ca^2+^ oscillations (de Vrind et al., 2016; Ping and Shepard, 1996; Takakusaki and Kitai, 1997). In PPN neurons, nifedipine could be blocking the high-threshold L-type Ca^2+^ channels activated during the AP spike, leading to less Ca^2+^ activation of SK channels to deepen the afterhyperpolarization trough and interspike Vm. In SNc neurons, the unaltered interspike Vm after nifedipine treatment could be the result of changing multiple conductances that compensate one another’s effect. While the low-threshold L-type Ca^2+^ channels are blocked by nifedipine, which would reduce interspike subthreshold depolarization, there is less Ca^2+^ entry to activate SK channels and SK-mediated hyperpolarization. Therefore, our results from studying AP kinetics are consistent with the idea that SNc neurons predominantly express the low-threshold subtypes and PPN neurons the high-threshold subtypes of L-type Ca^2+^ channels.

Interestingly, we observed heterogeneity within PPN cholinergic neuron responses to nifedipine treatment. While nifedipine did not have a statistically significant effect on the levels of tonic Ca^2+^ in PPN cholinergic neurons as a whole, we found that nifedipine decreased the firing frequencies and tonic Ca^2+^ levels while increasing pacemaking irregularity in almost half of the PPN neurons. The firing frequencies and irregularity in the rest of the PPN neurons were unchanged or increased after nifedipine. This suggests the possibility that a subpopulation of PPN neurons may rely on L-type channels for pacemaking and tonic Ca^2+^. PPN cholinergic neurons have historically been divided into several subgroups based on their electrophysiological properties and anatomical locations (Baksa et al., 2019; Kang and Kitai, 1990). Future studies are needed to determine whether dendritic Ca^2+^ signaling in PPN neurons differs in specific subpopulations.

In summary, our study shows that PPN cholinergic neurons do not share the characteristic of having low-threshold L-type Ca^2+^ conductance with SNc dopaminergic neurons, and thus there are likely other factors that underlie the selective vulnerability of cholinergic PPN neurons to degeneration. The lack of subthreshold Ca^2+^ conductance may be related to clinical observations that PPN neurons (∼30-60% loss) do not degenerate to the same extent as SNc neurons (∼70% loss) in PD, and PPN neurons have a larger between-patient variation in the percentage loss (Giguère et al., 2018). Many other intrinsic factors could also contribute to cellular vulnerability. Those relevant to PPN neurons include spontaneous pacemaking activity, having a large soma and extensive axonal arbor, dysfunctional proteostasis, and mitochondrial oxidative stress. Here we show that PPN neurons exhibit spontaneous firing activity and significant tonic Ca^2+^ entry, even though the main source of this pacemaking Ca^2+^ is not low-threshold L-type Ca^2+^ channels. Our data also show that PPN neurons and SNc neurons have comparable soma size and membrane capacitance, suggesting they likely have similar morphology and bioenergetic burden. Future work is needed to determine whether PPN cholinergic neurons are prone to the same proteostatic and mitochondrial stress as SNc dopaminergic neurons, and whether other Ca^2+^ channel subtypes contribute to the vulnerability of brainstem neurons such as the PPN.

## Acknowledgements

This project was funded by NIH BRAIN Initiative grant R00NS112417, American Parkinson’s Disease Association (APDA) grant R13 2021APDA00RG00000209666, and Parkinson’s Disease Foundation Stanley Fahn Junior Faculty Award PF-SF-JFA-10400267 awarded to RCE. We would like to thank Chelsea Scott and the Georgetown University Department of Comparative Medicine for maintaining our animal colony.

