## Supplementary Fig for "Comparing tonic and phasic calcium in the dendrites of vulnerable midbrain neurons"

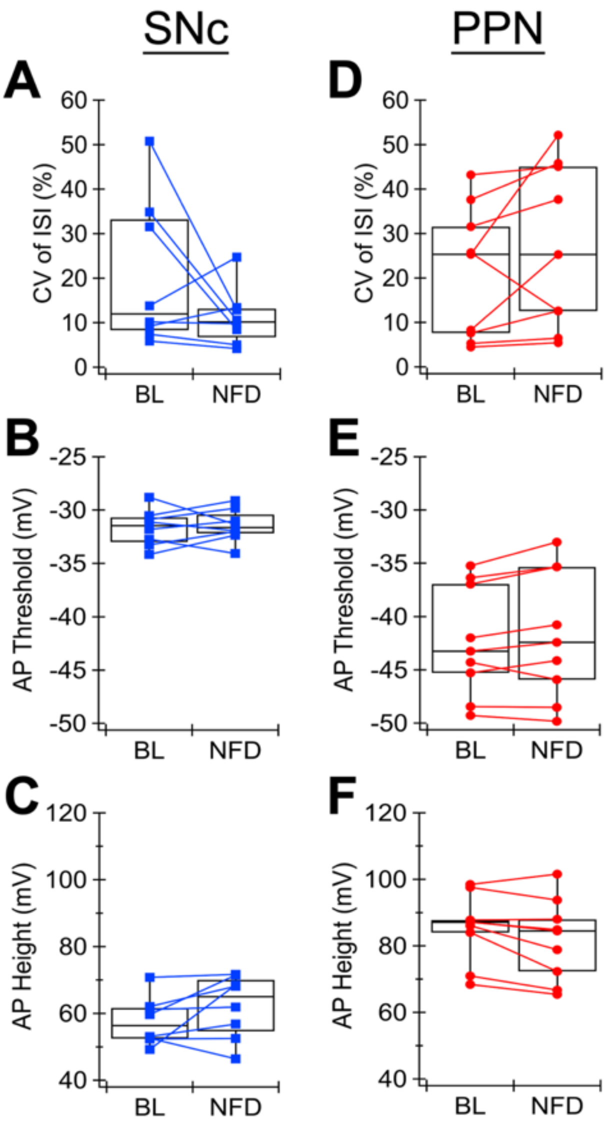


**Supplementary Figure 1. (A)** The firing regularity, represented by the coefficient of variation (CV) of the interspike interval (ISI), **(B)** AP threshold potential, and **(C)** AP spike height of SNc dopaminergic neurons at the baseline (BL) and after bath treatment of nifedipine (NFD, 10 μM). **(D)** The firing regularity, **(E)** AP threshold potential, and **(F)** AP spike height of PPN cholinergic neurons at the baseline and after nifedipine treatment.

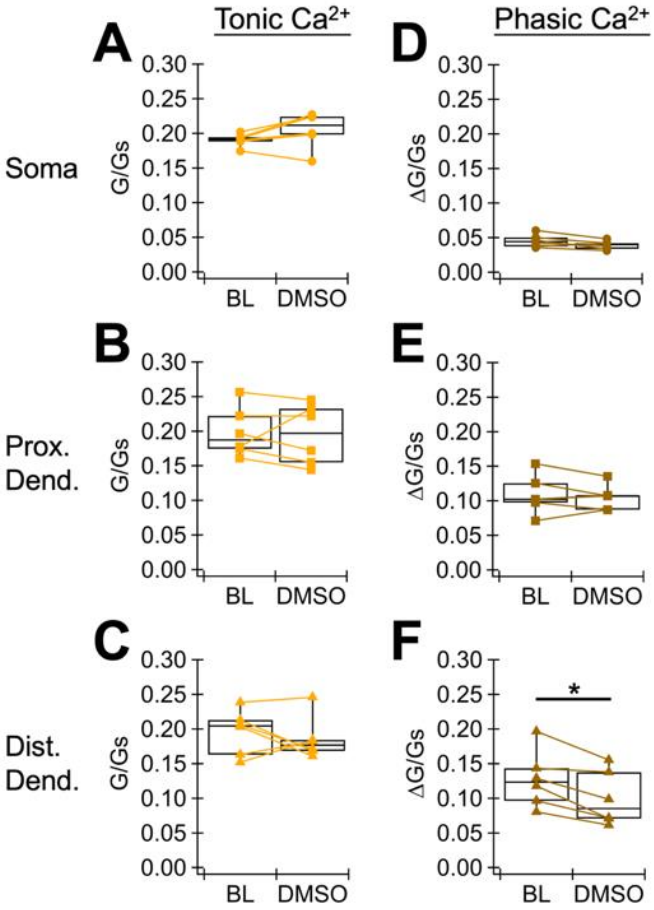


**Supplementary Figure 2. (A)** Summary box plot of the basal $Ca^{2+}$ levels (during tonic firing at 0 pA holding current) measured at the soma, **(B)** proximal dendrite, and **(C)** distal dendrite of PPN cholinergic neurons at the baseline (BL) and after bath treatment of DMSO (0.05%). **(D)** Summary box plot of the phasic $Ca^{2+}$ amplitudes (peak – basal $Ca^{2+}$) evoked by a 200-pA current step measured at the soma, **(E)** proximal dendrite, and **(F)** distal dendrite of PPN cholinergic neurons at the baseline and after DMSO treatment.
